# Demographic disequilibrium and growth-mortality asymmetry of forests across climate gradients

**DOI:** 10.1101/2023.06.25.546473

**Authors:** Jian Zhou

## Abstract

Forests are commonly believed to adapt themselves to environment and ultimately converge to demographic equilibrium characterized by a fixed size-structure. The expectation, however, has been lacking in mathematical rigor and been debating for evidential solidity. Here, by giving a general time-dynamic solution to the forest demographic model and verifying the prediction with worldwide forest inventory data, we show the inherent disequilibrium of forest demography with oscillations of forest-size-structure. Forests adapt to environment in a way of asymmetric growth-mortality tradeoff along climate gradients, which generates divergence to convergence oscillations of size-structure with rising temperature and precipitation. The demographic disequilibrium framework can provide a general basis for elucidating the variability of forest-size-structure with implications on intrinsic ecosystem instability, and for improving the Earth system modeling.

## Main Text

Forests play a vital role in the global eco- and climate-system for carbon sequestration and thus climate mitigation, and meanwhile, are highly impacted by climate change(*1, 2*). Forests adaptation to climate and their carbon sink potential are typically studied in two aspects: climate effects on physiologic processes (e.g., photosynthesis and respiration) and on demographic processes (e.g., growth and mortality)(*3-5*). The former are better understood as they act at shorter timescales and have been extensively studied with models and experiments, whereas the latter act at longer time scales and usually exhibit uncoupling with the former(*4*), making them the main source of uncertainty in forest performances. Although forest demography has long been investigated and is getting increasingly concerned in Earth system models in recent years(*3, 6-9*), our knowledges are still quite limited in the laws governing tree growth and mortality along climate gradients and their consequences on forest structure dynamics.

Following the conception of scale-independence in biology, early studies have attempted to scale up tree metabolic attributes to forest demographic properties with consistency in principles(*10, 11*). By applying the rule of energetic equivalence [total metabolic rates and corresponding total resource use keep constant during size-class transition(*12*)] to constraint the relationship between tree growth and mortality, the metabolic theory of ecology proposes an invariant scaling relation between tree size and abundance in natural forests across the globe(*13*). This is suggested to be consistent with the scaling relation between branch size and abundance of a tree(*14*). Despite the support of some combined data from multiple forest transects(*10*), many inventory data from large forest plots, however, contradict this idea(*15, 16*). The theory has also been heavily contested due to its lack of consideration of asymmetric competition for light(*17*), and mathematical problems in its scaling up(*18*).

Putting aside the pursuit of the consistent scaling rule in forests, modern forest demographic theories have been based on the more generalized mathematical analysis of the relationship between tree growth (*G*), mortality (*μ*), recruitment (*ρ*), and forest size structure (FSS) dynamics(*15, 19, 20*). Theoretically, given an initial FSS, the change of tree number (*N*) in each size-class (*S*) with time (*t*) can be estimated as the sum of the number of trees entering and leaving the size-class due to growth minus the number of trees dying in that size-class over time. Recruitment occurs only in the smallest size class (*S*_0_) by the establishment of seedlings or propagules in natural forests(*21*). Typically, a partial differential equation (PDE) model is used to provide the mathematical description:

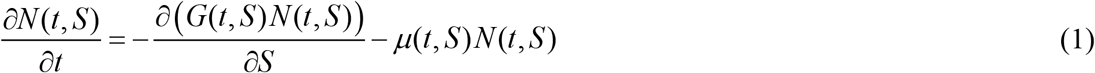

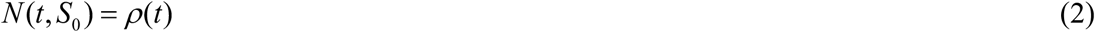

Despite of the simplicity in the functional form, finding a general time-dynamic solution to the model has been a tricky problem. As a result, our ability to comprehend dynamic FSS and forest carbon sequestration has been severely hampered(*21*). For decades, most of the relevant work has been based on the pre-assumed demographic equilibrium state and a static solution to the model (i.e., letting ∂*N*/∂*t* = 0). In this sense, all it does is to show how distinct patterns of FSS at the steady state might result from shifts in the size-specific growth and death rates(*15, 22, 23*). However, it is still uncertain whether forests will inherently tend towards the demographic equilibrium from any initial states and under any boundary conditions, and how forest demographic properties and FSS react to environmental changes. Addressing both issues requires developing a novel approach for analyzing FSS’s time-dynamic behavior and taking into account the possibility that adaptations to shifting environmental conditions would result in different dynamic rather than static FSS patterns.

In mathematics, difficulties in solving the PDE model derive from the definition on the boundary condition (i.e., saplings recruitment). Current studies of forest dynamic modeling usually define recruitment as a function of reproduction with detailed considerations on the age-or size-specific reproductivity of different species(*8, 9, 21, 24*). This introduces a great deal of complexity to the model solution. While the definition is more applicable to animal populations, in which reproductivity is the primary driver of population structure dynamics(*25*), seeds and propagules in plant communities are generally abundant in natural habitats irrespective of species composition(*26*). As such, saplings can recruit as far as essential environmental resources (e.g., space, light, and water) are sufficient and available(*27*). Recruitment in a natural forest could be thus defined as a resource-dependent rather than reproduction-dependent process. With instant sapling recruitment, there should be a constant equilibrium between the overall occupation of the limiting resource and the resource supply in a forest, which can be expressed as:

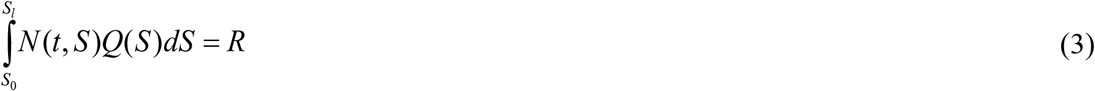

where *Q*(*S*) represents the size-dependent resource occupation capacity of a tree; *S*_0_ and *S*_*l*_ are the smallest (i.e., saplings) and the largest size of trees in a forest; *R* represents the total limiting resource for tree establishment in the forest and can be considered as a constant under a certain environmental condition.

To facilitate mathematical derivations of the model with the new definition of the boundary condition, we define *N*(*t, S*) as a multiplication of two parts based on the principle of variable separation(*28*):

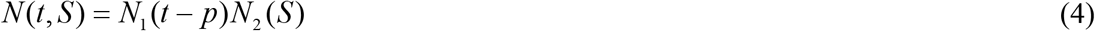

where *N*_1_(*t* – *p*) represents the normalized time-dynamic function of each size class initiated from the recruitment dynamics (*p* is the phase difference between the time-dynamics of size-class ‘*S*’ and the smallest size-class ‘*S*_0_’); *N*_2_(*S*) represents the size-dependent function of tree density that determined by growth and mortality, which equals to the static solution.

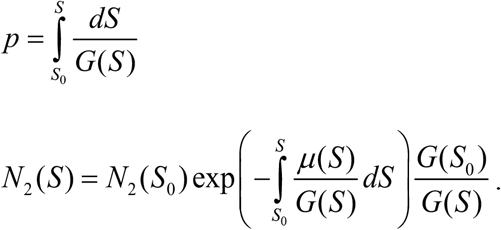

Then we introduce a function of *Q*(*S*) based on the fact that in a crowded competitive plant population, the increase of individual resource occupation with plant growth is at the expense of plant mortality (*18*), which can be expressed as:

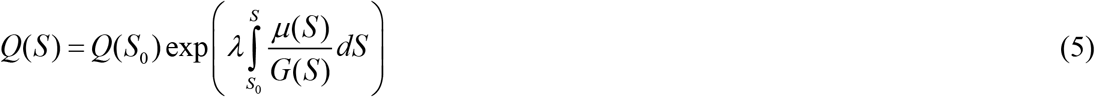

where *λ* is a parameter that regulates the relationship between the actual size-dependent resource occupation and the size-specific mortality to growth ratio of trees.

In general, the relationship between resource occupation and tree size is relatively conservative as it is fundamentally constrained by the geometric structure of vascular plants, whereas the size-specific mortality and growth are typically variable in response to environmental conditions. Consequently, for the application of Eq. 5 in actual conditions, we may anticipate a changing value of *λ* as the environment changes (e.g., the increase of *μ*/*G* will lead to the decrease of *λ*).

Specifically, *λ* = 1 corresponds to the condition that resources released from mortality are just fully occupied by growth during the process of a size-class transition (corresponds to the condition of ‘energetic equivalence’); while *λ* < 1 and *λ* > 1 correspond to the conditions in which resources occupied by growth are less and greater than those released by mortality, respectively. Under the constraint of resource release-occupation balance of the forest, the change of *λ* will further relate to the resource sufficiency for saplings recruitment (Fig. 1).

**Fig. 1.**
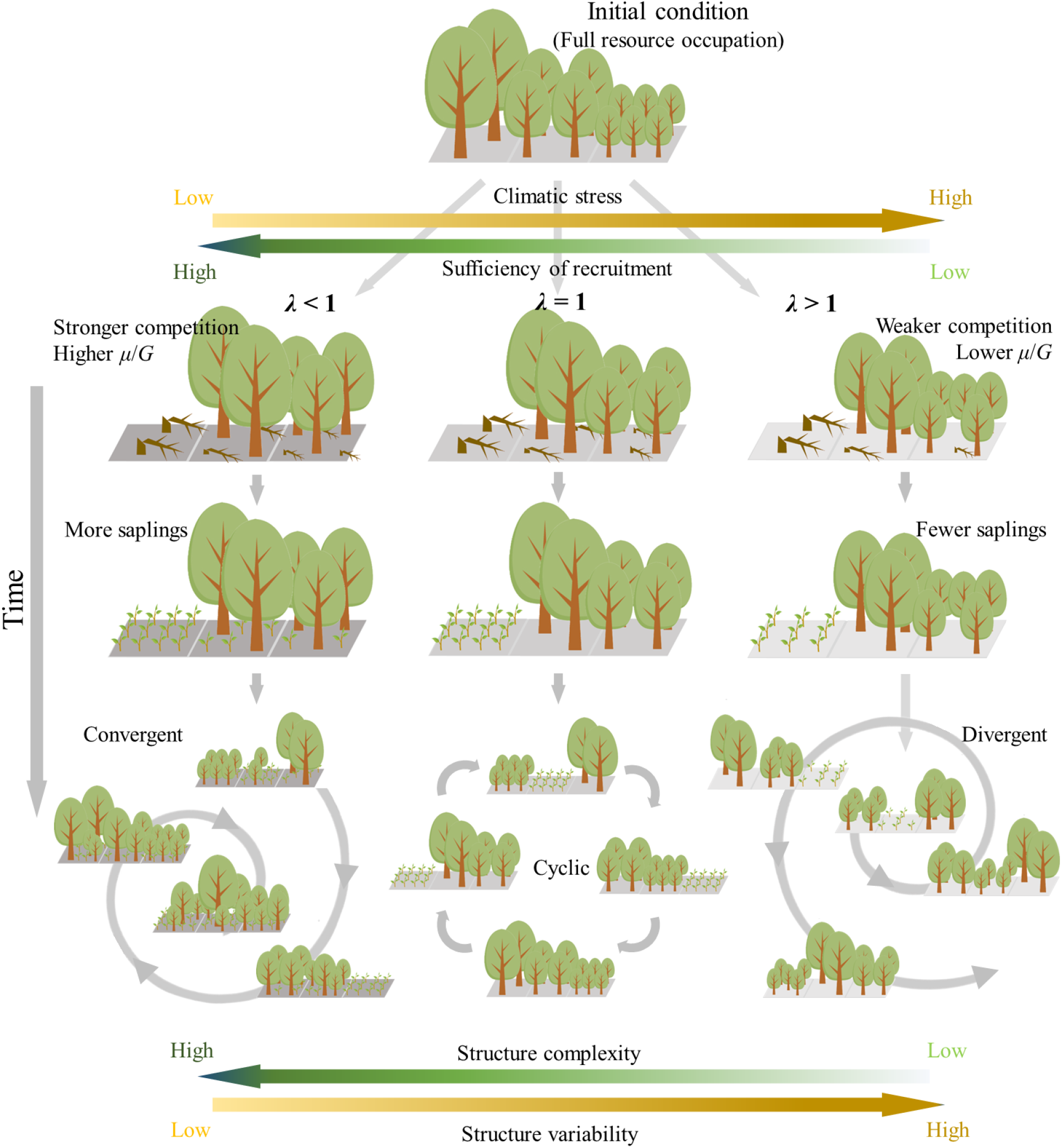
A schematic diagram of forest size structure dynamics under different scenarios. With increasing climatic stress, tree mortality to growth ratio and the associated recruitment decreases, leading to higher variability and lower complexity of forest size structure.

Putting Eqs. 4 and 5 into Eq. 3 results in a criterion of time-dynamic patterns of *N*_1_(*t*):

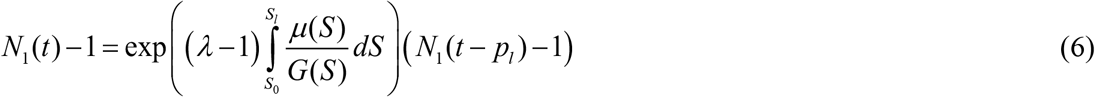

where *p*_*l*_ is the time spent for an individual growing from *S*_0_ to *S*_*l*_, i.e., the maximum lifespan of trees in the forest.

This finding demonstrates that the value of *λ* fundamentally determines the time-dynamic pattern of FSS. When *λ* = 1, *N*_1_(*t*) = *N*_1_(*t* – *p*_*l*_), which denotes a periodic function with the period length of the maximum tree lifespan. Under this condition, FSS will display a periodic oscillation if the forest is not initially at demographic equilibrium (Fig. 2A_2_). When *λ* < 1, *N*_1_ will converge to 1 with time, which agrees with the commonsense idea of FSS’s inherent convergence (Fig. 2A_1_). *λ* > 1 corresponds to a diverging function (Fig. 2A_3_). This will ultimately lead to a pulsing oscillation of FSS with only one size-class left in the forest as shown in subpanels in Fig. 2A_3 &_ B_3_.

**Fig. 2.**
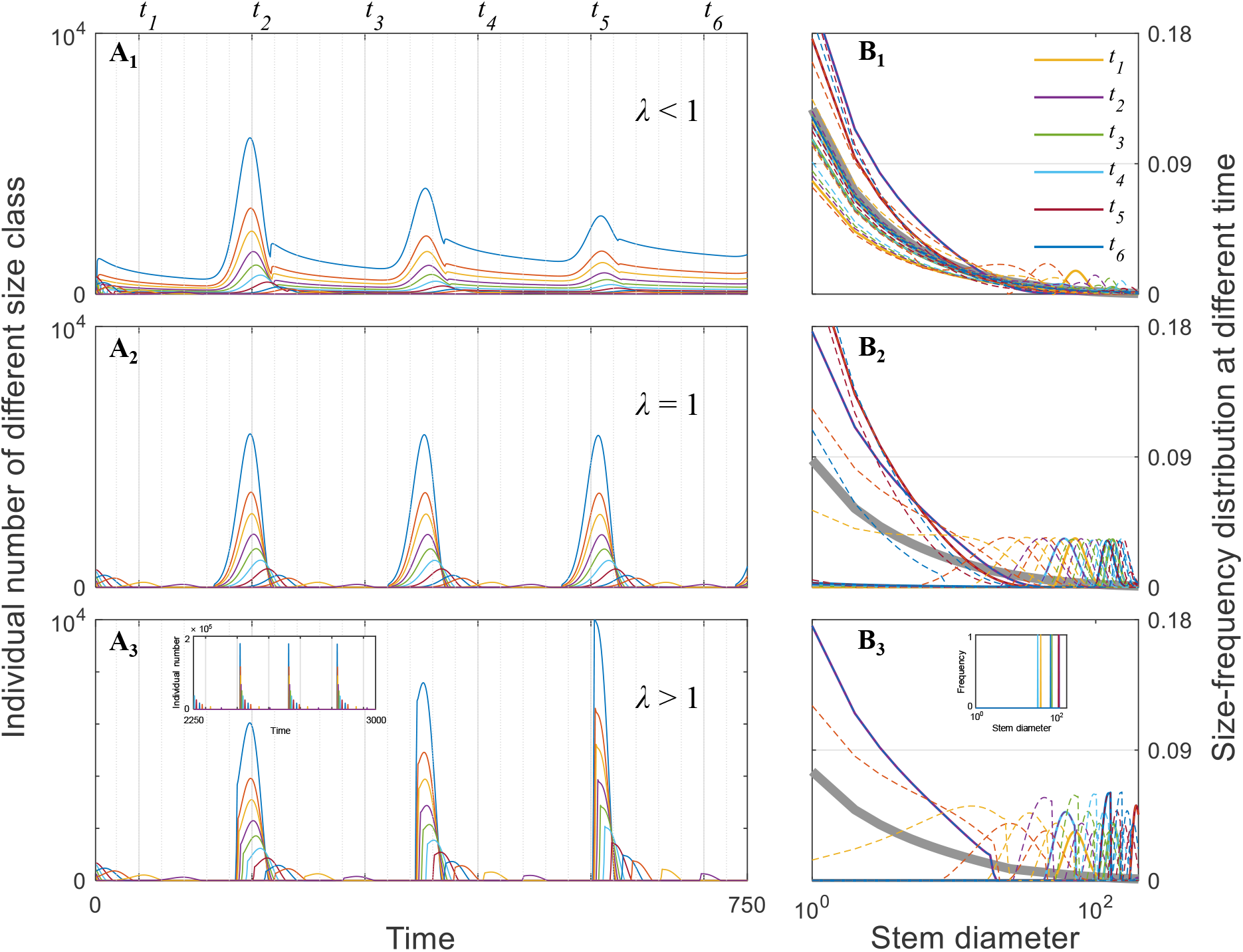
Simulated dynamical patterns of forest size structure under different value of *λ*. Colored lines in (**A**_**1**_ to **A**_**3**_) illustrate how tree numbers in different size-classes change over time. Colored lines in (**B**_**1**_ to **B**_**3**_) represent the tree size-frequency distribution (SFD) patterns at different times, which were marked with vertical lines in (**A**_**1**_ to **A**_**3**_), in which (***t***_**1**_ to ***t***_**6**_) were drawn with solid lines and others with dashed lines. The SFDs that merged at various dates are depicted as thick grey lines in (**B**_**1**_ to **B**_**3**_).

Determination of the value of *λ* in natural forests — according to our definition in Eq. 5 — requires an evaluation on the effect of environmental conditions on the size-specific *μ*/*G*. Using massive forest inventory data across the globe(*29-31*), we find that for the size-specific tree growth and mortality which estimated as *G* ∝ *S*^*α*^ and *μ* ∝ *S*^*β*^, the exponent of *μ*/*G* (i.e., *β* – *α*) shows an increasing trend with mean annual temperature (MAT) and precipitation (MAP) across the globe (Fig. 3B_2_ &C_2_). This finding implies that along climate gradients, the change in mortality is greater than the rate of change in growth. This represents an asymmetric growth-mortality tradeoff that break the hypothesis of universal ‘energetic equivalence’ (Fig. 3A). In consequence, the value of *λ* will tend to decrease with the increase of MAT and MAP (Fig. 3B_3_ & C_3_). Mechanisms for the asymmetric growth-mortality tradeoff needs in-depth study, but it can be basically inferred from the stress-gradient-hypothesis that tree competition (e.g., for light) tends to be stronger under more benign physical environmental conditions(*32*). Therefore, the stronger competition could lead to a greater increase in mortality along resource stress gradients.

**Fig. 3.**
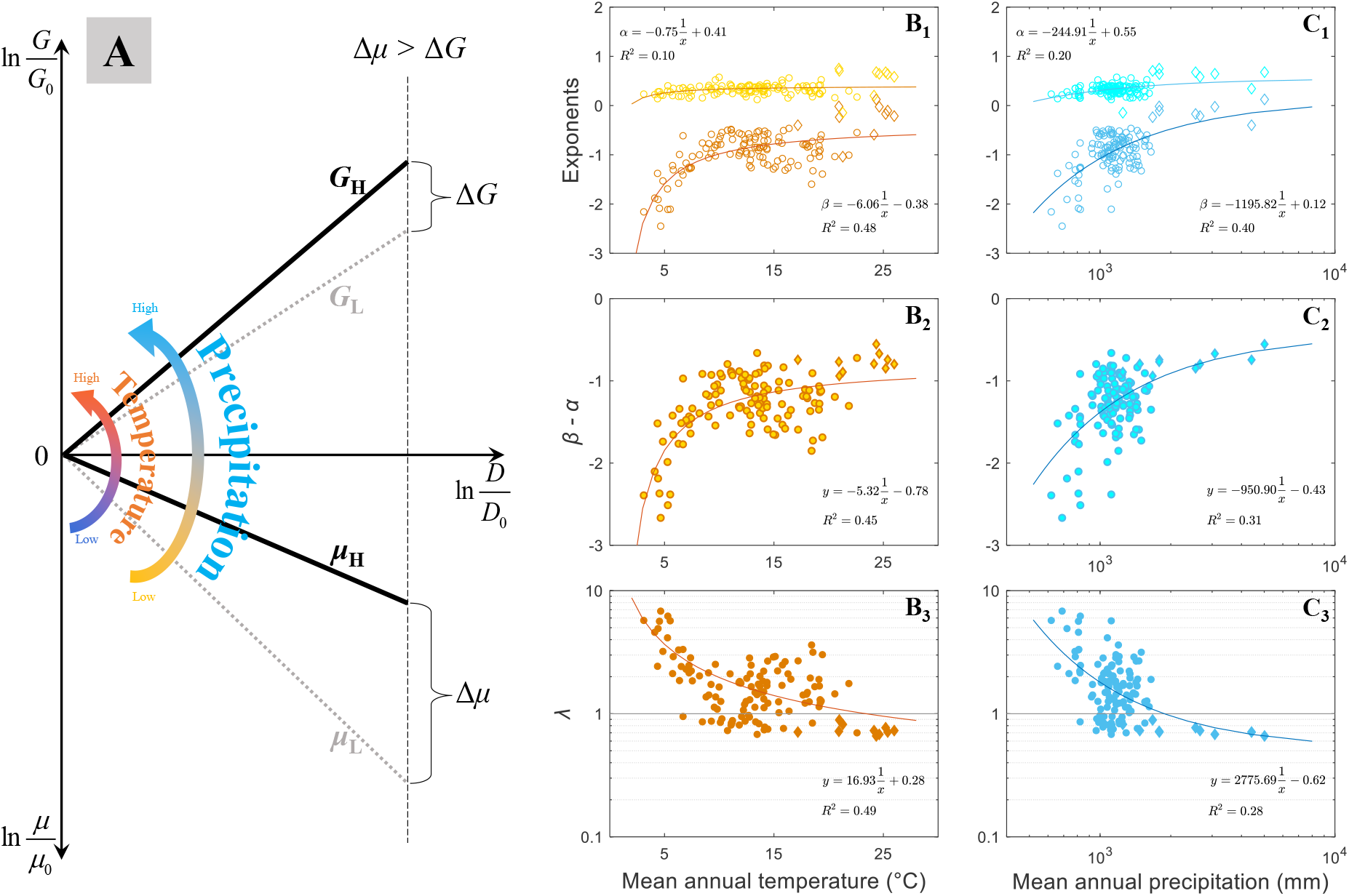
Asymmetric forest growth-mortality tradeoff along climate gradients. (**A**) A conceptual representation of the asymmetry. *G*_L_, *G*_H_ and *μ*_L_, *μ*_H_ represent size-specific growth and mortality rates under low and high temperature and precipitation, respectively. (**B**_**1**,**2**,**3**_ and **C**_**1**,**2**,**3**_) Changes of the growth and mortality exponents and the value of *λ* along temperature and precipitation gradients. Estimated data from the Forest Inventory Analysis (FIA) program and the Large-scale Tropical Forest Plots Network are shown in the circle and diamond plots, respectively.

With the determination of the effect of climates on the exponent of *μ*/*G* as well as the value of *λ*, we can predict that for *N*(*t, S*) = *N*_1_(*t* – *p*)*N*_2_(*S*), with precipitation and temperature rise, the slope of *N*_2_(*S*) will get steeper; while with precipitation and temperature fall, the oscillation of *N*_1_(*t* – *p*) will get stronger. For verification, we take two indirect indices to characterize the variation pattern of FSS along climate gradients: (i) the average of the size distribution exponent (DEX-av) and, (ii) the coefficient of variation of the size distribution exponent (DEX-cv) of forest inventory plots in each climate gradient. It follows that the steeper slope of *N*_2_(*S*) under higher MAT and MAP should correspond to the higher value of DEX-av, while the stronger oscillatory of *N*_1_(*t* – *p*) under lower MAT and MAP should correspond to the higher value of DEX-cv (Fig. 4A-B). Direct portraits of the FSS of old-growth natural forests along precipitation gradient (Fig. 4C_1_-C_3_) also indicate great consistency with the prediction (Fig. 2B_1_-B_3_). With decreasing precipitation and increasing value of *λ*, the size distributions show increasingly divergence in the head and more peaks in the tail. This corresponds to a greater number of phases in forest succession that are demographically out of balance.

**Fig. 4.**
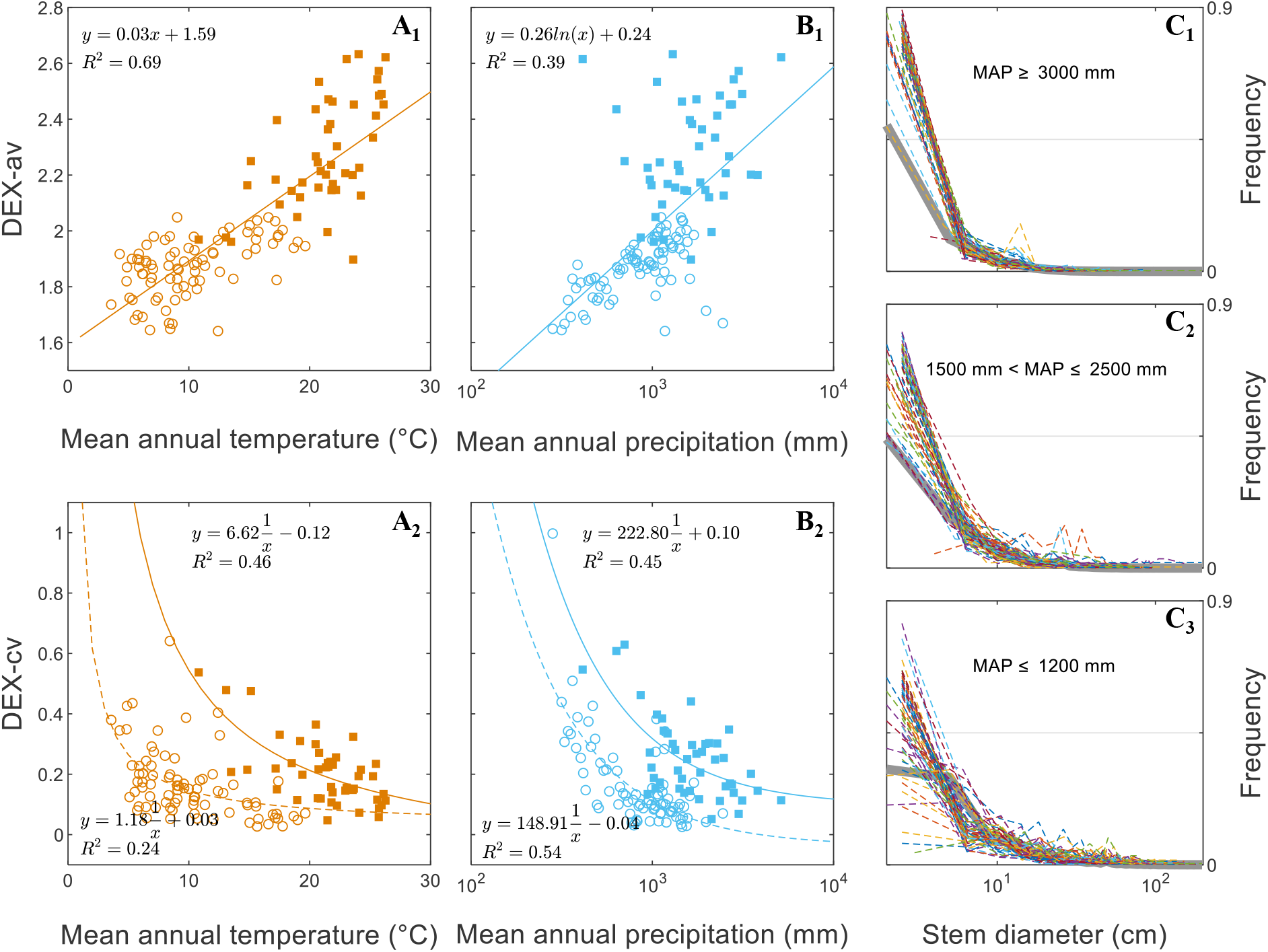
Variability patterns of forest size structure along climate gradients. (**A**_**1**,**2**_ and **B**_**1**,**2**_) The increase of DEX-av and the decrease of DEX-cv with the increase of MAT and MAP. Points shown by filled squares and empty circles were estimated using Gentry’s Forest Transect data and grid scale FIA data, respectively. The larger spatial scale of FIA data leads to the relatively lower DEX-cv in **A**_**2**_ and **B**_**2**_. (**C**_**1**,**2**,**3**_) Direct portrait of the variation of SFD patterns across precipitation gradients with Gentry data (60 transects in each precipitation gradient). The SFD of a transect is depicted by each colored dashed line. Each thick grey line represents SFD of the combined data of the 60 transects in each precipitation gradient.

In statistics, a combination of many of the transect or quadrat scale forest inventory data usually leads to a relatively stable FSS which is usually taken to represent approximately the demographic equilibrium state(*33*) (thick grey lines in Fig. 4C). This corresponds to the concept of shifting-mosaic steady state in which the random asynchrony of local scale patch dynamics could lead to a relatively stable state of the regional scale landscape (*34*). But strictly speaking, this regional equilibrium concept requires the cyclic dynamic pattern of local forests, which corresponds to the condition of *λ* = 1 in our mathematical framework. In climate zones where *λ* ≠ 1, the convergent or divergent oscillation of FSS cannot be totally smoothed out even under completely random spatial asynchrony. This can be mathematically demonstrated: the integral of a periodic function over a whole period will result in a time-independent constant, but the same integral of divergent or convergent oscillation functions will still be time-dependent. Spatial asynchrony does help in weakening the oscillation amplitude of FSS but cannot completely eliminate it (Fig. 4 A_2_ and B_2_ show the lower DEX-cv in a larger spatial area). In this sense, demographic disequilibrium is arguably a general intrinsic attribute of forests from local to global scales.

Our study provides both a theoretical framework and empirical evidence to demonstrate how interactions between growth, mortality and recruitment under different climate conditions lead to different dynamic patterns of FSS. We show that demographic equilibrium is not always a natural trend in forest succession, especially in regions where climatic stress is strong and recruitment is insufficient. The intrinsic oscillations of FSS implicates oscillations in forest carbon stocks and thus intrinsic disequilibrium of forest carbon dynamics. As such, the instability of ecosystems does not necessarily derive from external disturbances or climate changes. Rather, small random disturbances are essential for the ecosystem stability maintenance at large spatial scales as they facilitate spatial asynchrony. Moreover, climate changes do not simply drive shifts in the static equilibrium points of ecosystem states as previously concerned, but will lead to the change of dynamic disequilibrium patterns. Collectively, our results highlight the importance of accounting for the ecosystem’s intrinsic demographic interactions and disequilibrium with implications for improving Earth system models in a rapidly changing climate.

## References and Notes

1. G. B. Bonan, Forests and climate change: forcings, feedbacks, and the climate benefits of forests. Science 320, 1444–1449 (2008). https://doi.org/10.1126/science.1155121

2. W. R. L. Anderegg et al., A climate risk analysis of Earth’s forests in the 21st century. Science 377, 1099–1103 (2022). https://doi.org/doi:10.1126/science.abp9723

3. N. G. McDowell et al., Pervasive shifts in forest dynamics in a changing world. Science 368, eaaz9463 (2020). https://doi.org/doi:10.1126/science.aaz9463

4. A. Cabon et al., Cross-biome synthesis of source versus sink limits to tree growth. Science 376, 758–761 (2022). https://doi.org/doi:10.1126/science.abm4875

5. J. K. Green, T. F. Keenan, The limits of forest carbon sequestration. Science 376, 692–693 (2022). https://doi.org/doi:10.1126/science.abo6547

6. N. Rüger et al., Demographic trade-offs predict tropical forest dynamics. Science 368, 165–168 (2020).https://doi.org/10.1126/science.aaz4797

7. R. A. Fisher, C. D. Koven, Perspectives on the future of land surface models and the challenges of representing complex terrestrial systems. Journal of Advances in Modeling Earth Systems 12, e2018MS001453 (2020). https://doi.org/https://doi.org/10.1029/2018MS001453

8. I. Martínez Cano et al., Allometric constraints and competition enable the simulation of size structure and carbon fluxes in a dynamic vegetation model of tropical forests (LM3PPA-TV). Global Change Biology 26, 4478–4494 (2020). https://doi.org/https://doi.org/10.1111/gcb.15188

9. E. Weng et al., Modeling demographic-driven vegetation dynamics and ecosystem biogeochemical cycling in NASA GISS’s Earth system model (ModelE-BiomeE v.1.0). Geosci. Model Dev. 15, 8153–8180 (2022).https://doi.org/10.5194/gmd-15-8153-2022

10. B. J. Enquist et al., Scaling metabolism from organisms to ecosystems. Nature 423, 639–642 (2003). https://doi.org/10.1038/nature01671

11. J. H. Brown, J. F. Gillooly, A. P. Allen, V. M. Savage, G. B. West, Toward a metabolic theory of ecology. Ecology 85, 1771–1789 (2004). https://doi.org/10.1890/03-9000

12. E. P. White, S. K. Ernest, A. J. Kerkhoff, B. J. Enquist, Relationships between body size and abundance in ecology. Trends in Ecology & Evolution 22, 323–330 (2007). https://doi.org/10.1016/j.tree.2007.03.007

13. B. J. Enquist, K. J. Nicklas, Invariant scaling relations across tree-dominated communities. Nature 410, 655–660 (2001). https://doi.org/10.1038/35070500

14. G. B. West, B. J. Enquist, J. H. Brown, A general quantitative theory of forest structure and dynamics. Proceedings of the National Academy of Sciences of the United States of America 106, 7040–7045 (2009). https://doi.org/10.1073/pnas.0812294106

15. H. C. Muller-Landau et al., Testing metabolic ecology theory for allometric scaling of tree size, growth and mortality in tropical forests. Ecology Letters 9, 575–588 (2006). https://doi.org/10.1111/j.1461-0248.2006.00904.x

16. H. C. Muller-Landau et al., Comparing tropical forest tree size distributions with the predictions of metabolic ecology and equilibrium models. Ecology Letters 9, 589–602 (2006). https://doi.org/10.1111/j.1461-0248.2006.00915.x

17. D. A. Coomes, R. B. Allen, Mortality and tree-size distributions in natural mixed-age forests. Journal of Ecology 95, 27–40 (2007). https://doi.org/10.1111/j.1365-2745.2006.01179.x

18. J. Zhou, G. Lin, Will forest size structure follow the -2 power-law distribution under ideal demographic equilibrium state? Journal of Theoretical Biology 452, 17–21 (2018). https://doi.org/10.1016/j.jtbi.2018.05.011

19. T. Hara, Dynamics of size structure in plant populations. Trends in Ecology & Evolution 3, 129–133 (1988). https://doi.org/10.1016/0169-5347(88)90175-9

20. R. Condit, R. Sukumar, Predicting population trends from size distributions: a direct test in a tropical tree community. The American Naturalist 152, 495–509 (1998). https://doi.org/10.1086/286186

21. P. R. Moorcroft, G. C. Hurtt, S. W. Pacala, A Method for scaling vegetation dynamics: the ecosystem demography model (ED). Ecological Monographs 71, 557–585 (2001). https://doi.org/10.1890/0012-9615(2001)071[0557:AMFSVD]2.0.CO;2

22. C. E. Farrior, S. A. Bohlman, S. Hubbell, S. W. Pacala, Dominance of the suppressed: Power-law size structure in tropical forests. Science 351, 155–157 (2016). https://doi.org/10.1126/science.aad0592

23. J. R. Moore, A. P. K. Argles, K. Zhu, C. Huntingford, P. M. Cox, Validation of demographic equilibrium theory against tree-size distributions and biomass density in Amazonia. Biogeosciences 17, 1013–1032 (2020). https://doi.org/10.5194/bg-17-1013-2020

24. D. Medvigy, S. Wofsy, J. Munger, D. Hollinger, P. Moorcroft, Mechanistic scaling of ecosystem function and dynamics in space and time: Ecosystem Demography model version 2. Journal of Geophysical Research: Biogeosciences 114, (2009). https://doi.org/10.1029/2008JG000812

25. B. Ebenman, L. Persson, Eds., Size-Structured Populations: Ecology and Evolution. (Springer-Verlag, 1988).

26. A. M. V. R. R. Warrier, C. Kunhikannan, Significance of soil seed bank in forest vegetation: a review. Seeds 1, 181–197 (2022). https://doi.org/10.3390/seeds1030016

27. T. Valverde, J. Silvertown, Canopy closure rate and forest structure. Ecology 78, 1555–1562 (1997). https://doi.org/10.2307/2266148

28. G. B. Arfken, H. J. Weber. Mathematical Methods for Physicists. (American Association of Physics Teachers, 1999).

29. O. Phillips, J. S. Miller, J. S. Miller, Global Patterns of Plant Diversity: Alwyn H. Gentry’s Forest Transect Data Set. (Missouri Botanical Press, 2002).

30. E. C. Losos, E. G. Leigh, Tropical Forest Diversity and Dynamism: Findings from a Large-scale Plot Network. (University of Chicago Press, 2004).

31. A. Gray, T. Brandeis, J. Shaw, W. McWilliams, P. Miles, Forest inventory and analysis satabase of the United States of America (FIA). Vegetation Databases for the 21st Century. Biodiversity and Ecology 4, 225–231 (2012). https://doi.org/10.7809/b-e.00079

32. F. T. Maestre, R. M. Callaway, F. Valladares, C. J. Lortie, Refining the stress-gradient hypothesis for competition and facilitation in plant communities. Journal of Ecology 97, 199–205 (2009). https://doi.org/10.1111/j.1365-2745.2008.01476.x

33. J. Westfall, Spatial-scale considerations for a large-area forest inventory regression model. Forestry 88, 267–274 (2015). https://doi.org/10.1093/forestry/cpv001

34. F. H. Bormann, G. E. Likens, Pattern and Process in a Forested Ecosystem: Disturbance, Development and the Steady State Based on the Hubbard Brook Ecosystem Study. (Springer-Verlag, 1994).

